# Novel gut microbiota and microbiota-metabolites signatures in gliomas and its predictive/prognosis functions

**DOI:** 10.1101/2023.01.19.524836

**Authors:** Min Zhou, Chong Song, Junwei Gu, Tong Wang, Linyong Shi, Chiyang Li, Liwen Zhu, Hong Li, Songtao Qi, Yuntao Lu

## Abstract

Gliomas are the most common malignant tumors in the central nervous system. Host genetic and environmental factors have been implicated as the causes and regulators of gliomas. Evidence shows that alterations of the gut microbiome play an important role in multiple diseases including central nervous system disorders. However, the influence of gut microbiome to the epigenesis of gliomas remains largely unknown. Here we profiled the gut microbiome and metabolome in fecal samples from healthy volunteers and the patients with gliomas through the 16S rRNA gene sequencing and LC-MS analyses. The fecal samples from primary glioma patients (n=51), recurrent glioma patients (n=11), patients who underwent TMZ radio-chemotherapy (n=16) and healthy volunteers (n=37) were collected. 56 discriminatory OTUs and 144 metabolites were observed in gliomas compared to those in healthy volunteers, and some species were correlated with clinical parameters, such as tumor grade, IDH-1 and MGMT status. Moreover, the gliomas group showed increased activity in pathways associated with ectoine biosynthesis, fatty acid elongation (saturated), and protocatechuate degradation. At the same time, we revealed 4 fatty-acid metabolites(palmitic acid, oleic acid, DL−beta−Hydroxypalmitic acid, 4−(Methylamino)−4−(3−pyridyl)butyric acid) as possibly interacting with glioma growth. Random forest modeling indicated that a model involving 8 genera and 10 metabolite biomarkers achieved a high accuracy in gliomas prediction (AUC=94.4%). We investigated interassociations between the microbial genera in glioblastoma multiforme (GBM) and progression-free survival (PFS) and overall survival (OS) by Spearman’s correlation analysis. Patients with high proportions of fecal *Faecalibacterium* had significantly better median PFS or OS than those with low proportions (mPFS 495 vs. 281 days, p=0.005; mOS 604 vs. 395 days, p=0.044). Moreover, animal experiments have verified the causal relationship between the structural changes of gut microbiome and glioma growth. Our current study comprehensively characterizes the perturbed interface of gut microbiome and metabolites in glioma patients, which may be used as diagnostic and prognostic biomarkers of gliomas.

## Introduction

Gliomas are the most common primary malignant tumors of the central nervous system (CNS) in adults, accounting for approximately 50% of all the primary brain tumors(1). The current standard treatment of newly diagnosed gliomas is maximal surgical resection followed by radiotherapy combined with temozolomide (TMZ) chemotherapy(2). However, the prognosis is still relatively dismal, especially in patients with glioblastoma multiforme (GBM), whose median overall survival is 15 to 18 months(2, 3), To date, an increasing number of molecular markers have been identified that play key roles during glioma carcinogenesis and progression, but only a few of these biomarkers have been clinically translatable(4, 5). Thus, there is an urgent clinical need to develop noninvasive and more effective biomarkers for gliomas.

Increasing evidence has shown that both the composition and metabolism of the gut microbiota play an important role in not only gastrointestinal function but also systemic effects, exerting local and long-distance effects(6-8). Epidemiological and animal-based studies have suggested that the gut microbiota participates in the development and progression of cancer through mechanisms such as alteration in immune responses, and induction of genetic mutation(6, 9). Experimental evidence has shown that the gut microbiota is associated with lung cancer(10, 11), prostate cancer(12, 13), liver cancer(14) and other extraintestinal cancers(15). Naoki Sugimura et al(16) discovered that *Lactobacillus gallinaurm* and its metabolite (indole-3-lactic) protected against intestinal tumorigenesis. Perturbation of the gut microbiome,a pathologic disequilibrium, recently has been described in several CNS diseases such as multiple sclerosis(17), autism(18), Parkinson’s disease(19), and ischemic stroke(20). Ischemic stroke rapidly triggers gut dysbiosis with *Enterobacteriaceae* expansion and produces excessive nitrate, in turn exacerbates brain infarction(20). Hence, the term “brain-gut axis” is presented to describe the physiology between the CNS and the gastrointestinal system(21, 22). As one of the most common CNS malignant diseases, animal experimental studies have found that gliomas can induce the gut microbiome changes and are influenced by TMZ chemotherapy(23, 24). The microbiome is now considered as noninvasive biomarker for gliomas.

Few studies have reported on microbial associations with gliomas; thus, the profile of the gut microbiome of gliomas remains largely unknown. It is now clear that the metabolites of the gut microbiome, as biochemical converters, can exert tumor-promotive or tumor-suppressive functions(20, 25). An animal model study indicated metabolite changes in the feces of animal with gliomas(26). Nevertheless, the interplay between the gut microbiota and metabolites, and their roles in gliomas have not been effectively characterized. Therefore, we conducted this population study to analyze the gut microbiome of gliomas and healthy controls by sequencing the 16S ribosomal RNA (rRNA) gene sequencing and mass spectrometry. We assessed the gut microbial profiles and metabolite abundance in fecal samples from health volunteers and the patients with gliomas and identified the specific microbial and metabolic biomarkers of gliomas. Moreover, we deciphered the association of gliomas with the gut microbiota and fecal metabolites, and investigated the correlations between the gut microbiota and prognosis. Animal studies verified gliomas could influence compositions of gut microbiome. FMT (fecal microbiota transplantation) experiments certified high level of *Faecalibacterium* in gut could improve prognosis of gliomas.

## Results

### Clinical characteristics

The detailed clinical characteristics of the enrolled 37 healthy volunteers (N_Ctrl group) and 77 glioma patients are summarized in Table 1. The enrolled glioma patients contain three groups: patients with primary gliomas (Gliomas group), patients who underwent TMZ chemoradiotherapy (stupp regimen) (Treat group), and tumor recurrence after stupp treatment (Recurrence group). There were no significant differences in sex, age, or BMI among these groups. For the patients with primary WHO grade IV gliomas, we collected the follow-up information. The median PFS was 302 days (95% CI 257-347 days) and the median OS was 418 days (95% CI 364-471 days).

**Table 1.**
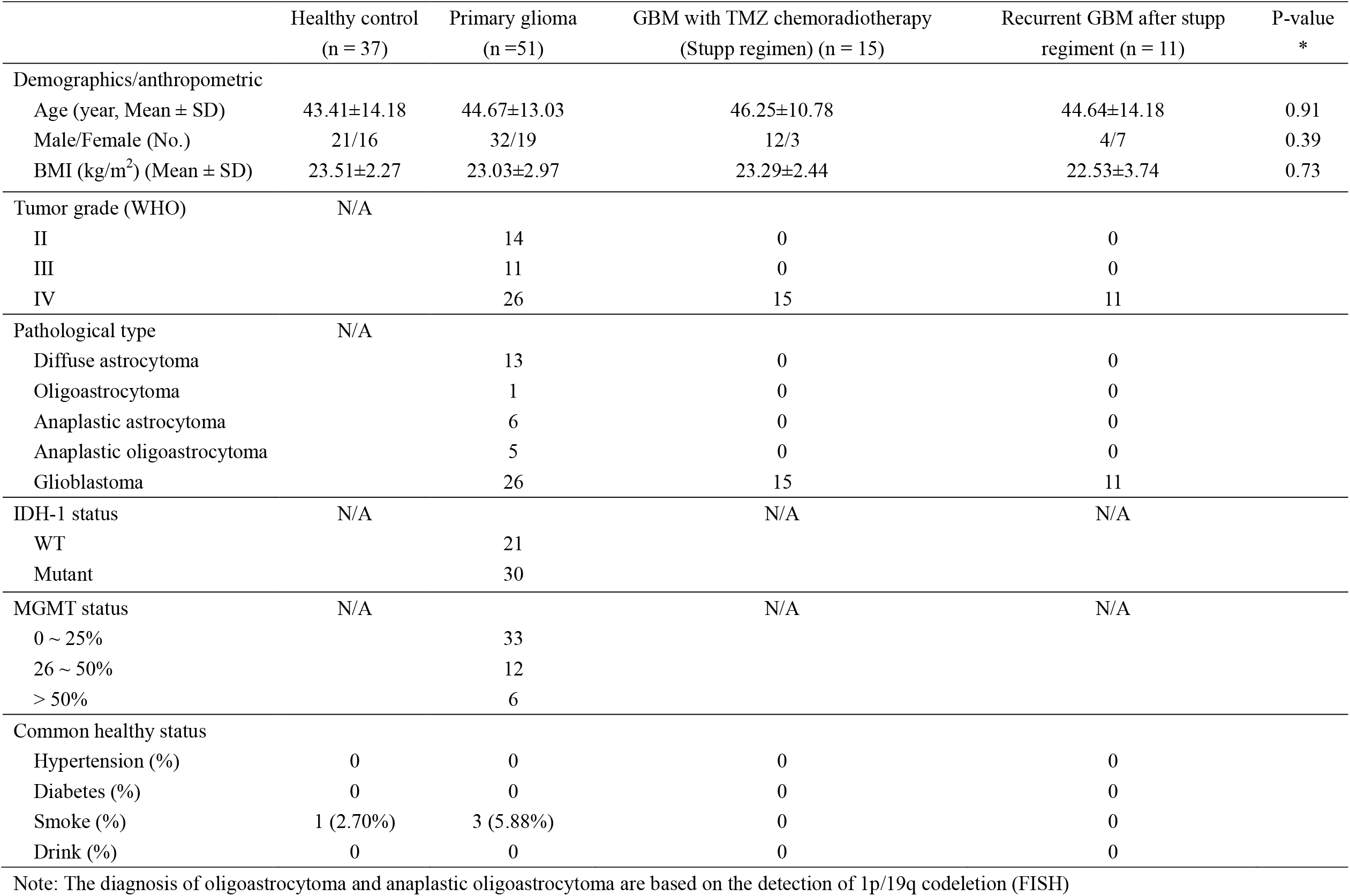
Baseline characteristics of our cohort.

### Alterations of the gut microbiota in primary glioma patients

From all the participant fecal samples, we obtained a total number of 5583731 high-quality reads, with a median read count of 48731 per sample. A total of 11714 features (as OTUs) were identified and labeled based on QIIME2 (Table S1). To assess the richness and diversity of the microbiota between the Gliomas group and the N_Ctrl group, we used the reads to evaluate alpha diversity and beta diversity. There were statistically significant differences in the Observed_OTUs (226.00±94.82 vs. 281.97±78.89, p=0.024) (Figure 1A) and Chao1 index (213.15±98.11 vs. 288.12±81.91, p=0.0087) (Figure S1A), whereas the Shannon index (4.80±1.09 vs. 5.06±0.93, p=0.14) (Figure 1B) and Simpson index (0.88±0.11 vs. 0.90±0.09, p=0.35) (Figure S1B) were not significantly different between the two groups. Principle coordinates analysis (PCoA) plot based on the unweighted UniFrac distances showed that the microbiome of the Gliomas group was different from that of the healthy control group. Analysis of similarities (ANOSIM) was performed, and the results revealed that the composition and abundance of the gut microbiome of the Gliomas group were significantly different from those of the healthy group (r=0.173, p=0.003, Figure 1C). These results indicated that the diversity of the gut microbiome may be strongly influenced during the development of gliomas.

**Figure 1.**
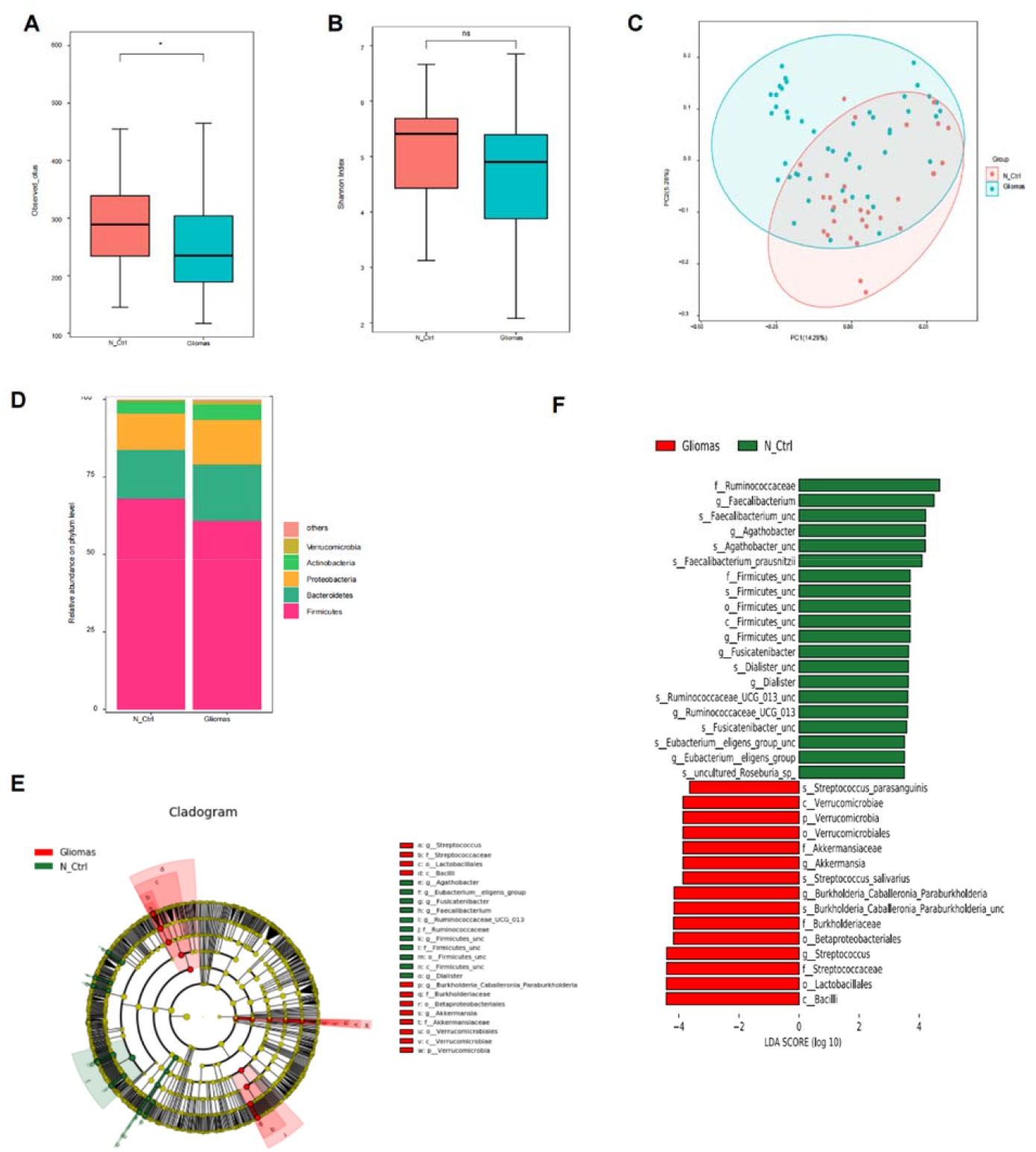
The shift of gut microbiota composition in gliomas patients. (A)(B) Microbial richness based on number of observed_outs, the shannon index. *p<0.05, **p<0.01, ***p<0.001. (C) Principal coordinates analysis (PCoA) for gliomas and N_Ctrl groups. Significant differences were observed between patients and controls with ADONIS test. The first two principal coordinates (PCs) were each labeled with the percentage of variance explained. (D) The relative abundance of bacterial phyla clustered into different groups was determined. (E) Cladograms generated by LEfSe indicating differences in the bacterial taxa between Gliomas and N_Ctrl (healthy controls) groups. Red bars indicate taxa were enrichment in Gliomas group, green bars indicate taxa were enrichment in N_Ctrl group. (F) LDA scores for the bacterial taxa differentially abundant between Gliomas and N-Ctrl groups (LDA > 3.0). Red bars indicate taxa were enrichment in Gliomas group, and green bars indicate taxa were enrichment in N_Ctrl group.

We compared the relative taxon abundances of the gut microbiome in Gliomas and healthy control groups. In total, 1003 microbial species representing 516 genera and 22 phyla were profiled in fecal samples of the two groups. At the phylum level, 14 phyla were identified in each group. *Candidatus Saccharibacteria, Planctomycetes, Deferribacteres, Gemmatimonadetes* were unique to gliomas. Compared to healthy controls, *Chloroflexi* and *Fibrobacteres* were absence in gliomas. In consistent with other gut microbiome studies, the prominent phyla in both groups were *Actinobacteria, Proteobacteria, Bacteroidetes* and *Firmicutes* (Figure 1D). Growing evidence shows that the alteration of *Firmicutes*/*Bacteroidetes* ratio (F/B) *is* an important indicator of gut microbiome imbalance. In this study, we compared the F/B ratio between two groups. Even though in Gliomas group the ratio decreased obviously, but there was no significant variation (p>0.05, Figure S1C). To further distinguish the predominant gut microbiota between the Gliomas and healthy control group, we used linear discriminant effect size (LEfSe) to identify the specific bacteria associated with gliomas (Figure 1E, 1F). We identified 57 key discriminatory OTUs (all LAD>3.0, Table S2). Among these, 3 phyla, 37 genera, 45 species showed significantly higher abundance in gliomas patients, whereas 1 phylum, 19 genera, 43 species were conversely depleted. *Verrucomicrobia*, a phylum comprising several important pathogens was much more enriched in patients with gliomas, compared to healthy controls. Several opportunistic pathogens including *Streptococcus, Burkholderia-Caballeroina-Paraburkholderia, Akkermansia, Citrobacter, Corynebacterium*, and *Blautia* were significantly overrepresented (all LDA scores >3.5) in the Gliomas group, whereas the microbiome at the genus level in the healthy control group was dominated by *Faecalibacterium, Agathobacter, Firmicutes, Fusicatenibacter, Dialister*, and *Ruminococcaceae_UCG-013* (all LDA scores >4.0) (p<0.05).

### Gut microbes correlate with glioma subtypes

We divided the primary gliomas groups into different subgroups according to IDH-1 or MGMT pathological status (Figure 2A and B), which is highly associated with malignancy and prognosis of gliomas. Spearman’s rank correlation coefficient method was used to evaluate the correlation between the gut microbiome and the pathological status. An IDH1-mutation status was significantly negatively correlated with the genera *Megamonas* (r=-0.45, p=0.030) and *Bacteroides* (r=-0.40, p=0.030), and positively correlated with the genera *Bifidobacterium* (r=0.38, p=0.043) and *Erysipelatoclostridium* (r=0.27, p=0.048). In addition, MGMT positive incidence was positively correlated with the genera *Collinsella* (r=0.35, p=0.029) and *Streptococcus* (r=0.40, p=0.030).

**Figure 2.**
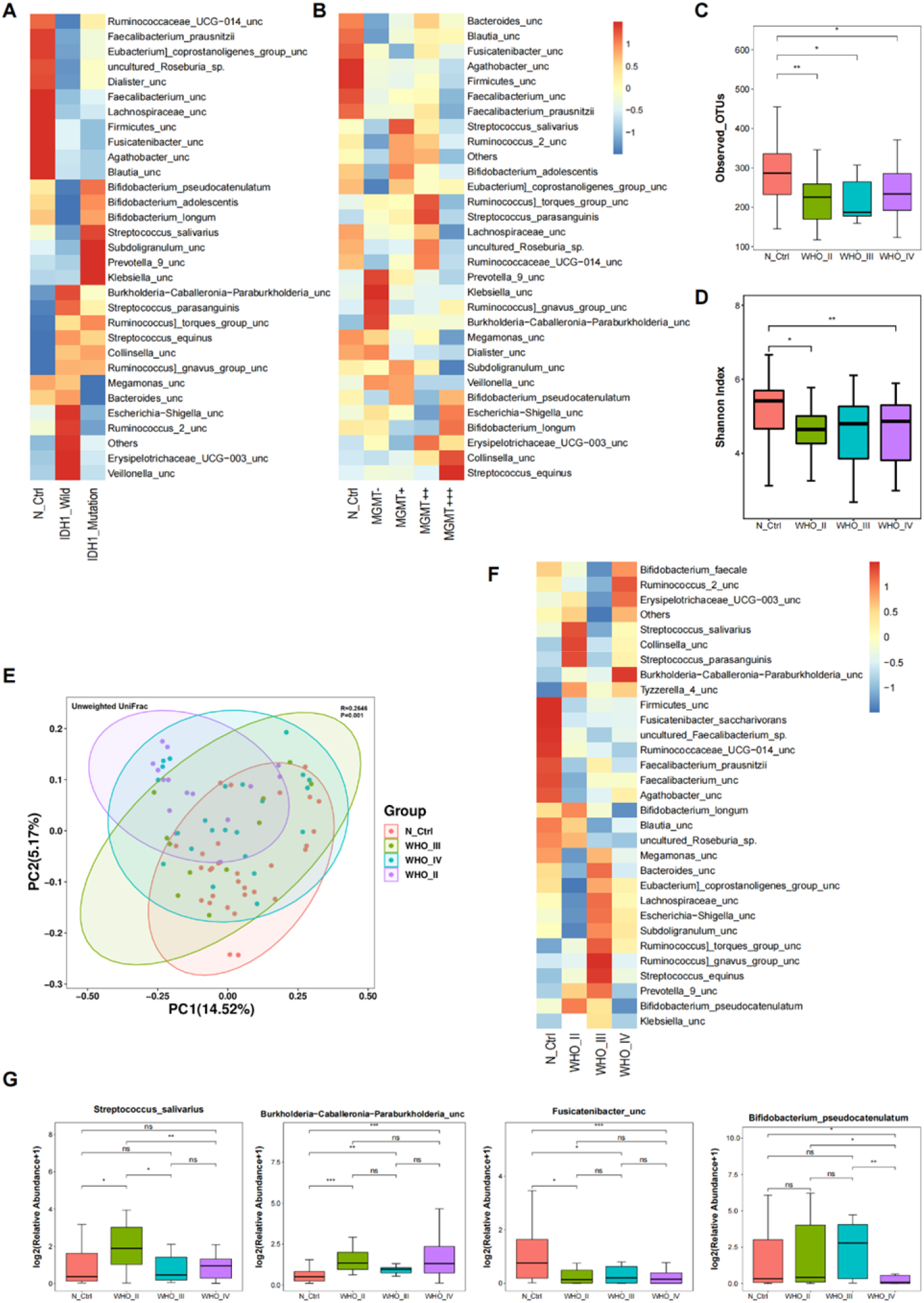
Association between clinical variables and gut microbiota abundance. **(A)(B)** Heat map showing the 30 most abundant species in samples grouped by IDH-1 and MGMT status. **(C)(D)** Microbial richness based on number of observed_otus, the shannon index. *p<0.05, **p<0.01, ***p<0.001. **(E)** Principal coordinates analysis (PCoA) for gliomas and N_Ctrl groups. Significant differences were observed between patients and controls with ADONIS test. The first two principal coordinates (PCs) were each labeled with the percentage of variance explained. (F) Heat map showing the relative abundances of the 30 most abundance species in samples grouped by pathological types. The relative abundance was indicated by color gradient. (G) The most significantly changed species were further analyzed in each group. Two-tailed Wilcoxon rank-sum test, *p <0.05, **p < 0.01, ***p < 0.001, ns: non-significance.

Next, according to WHO, we divided the gliomas group into three subgroups: the WHO_II group, III group, and IV groups. We compared the alpha and beta diversity in these three groups and the healthy control groups. There were statistically significant differences in the Observed_OTUs (p<0.001) (Figure 2C), but nonsignificant differences in the Shannon index (p=0.073) (Figure 2D). The analysis of beta diversity based on the unweighted UniFrac distances revealed that the microbiome of the three subgroups was significantly different from that of healthy control group (ANOSIM, r=0.265, p=0.001, Figure 2E). To confirm these differences, we compared the relative abundance at the species level (Figure 2F, G). These results showed a significant increase in the relative abundances of *Burkholderia-Calleroina-Paraburkholderia, Brevundimonas viscosa* and a significant decrease in *Citrobacter freundii, Fusicatenibacter, Citrobacter_sp, Fusicatenibacter, saccharovorans, Ruminococcaceae_UCG-013 and Clostridium_sp_AT4* in the three glioma subgroups compared to the healthy control group. Moreover, *Streptococcus_salivarius* significantly increased in WHO II subgroup, but *Bifidobacterium_pseudocatenulatum* decreased in WHO IV subgroup. We also performed LEfSe analysis (LDA>3.0, p<0.01) to identify the bacteria biomarkers between different grade gliomas. Compared to healthy controls, a similar reduction in three subgroups was found in the dominant phylum *Firmicutes*. With the increase of gliomas malignancy, we found enriched abundance of *Proteobacteria*, which could be explained by the increased abundance of Gammaproteobacteria, a class comprising several common opportunistic pathogens such as *Burkholderia-Calleroina-Paraburkholderia* and *Citrobacter*. All these results indicated that the constituents of the gut microbiome could be influenced by tumor grade. It is certainly necessary to recruit more patients for subtype-specific microbial studies in the future.

### KEGG pathways were altered in the gut microbiome of primary glioma patients

We further used the PICRUSt method to predict the KEGG pathways between the gut microbiome of primary glioma patients and healthy individuals. We found 157 different KEGG pathways at p<0.05. Among these pathways, 80 were increased and 77 were decreased in the Gliomas group (Table S3). The Gliomas group showed increased activity in pathways associated with ectoine biosynthesis, fatty acid elongation (saturated), and protocatechuate degradation. These results revealed the potential metabolic reprogramming in the gut microbiome during glioma development.

### Alteration of the gut microbiota in postoperative GBM patients with standard chemoradiotherapy and recurrent GBM patients

As GBM (WHO IV) is the most malignant pathological type of glioma with poorest prognosis, we conducted 16S rRNA gene sequencing to analyze fecal samples from postoperative glioma patients with standard therapy (Treat group) who received radiotherapy and four rounds of chemotherapy with TMZ, and patients suffered recurrent glioma after chemoradiotherapy (Recurrence group). The alpha and beta diversities were used to explore the variations in the gut microbiome among the healthy control group, primary GBM group, Treat group and Recurrence group. Comparison among these four groups, revealed significant differences in the Observed_OTUs (p=0.0048) and Shannon index (p=0.016) (Figure 3A and B). Then, we used the unweighted UniFrac method to analyze the beta diversity, and PCoA was performed. This indicated that there was a significant difference among the four groups (ANOSIM, r=0.224, p=0.04, Figure 3C). Thus, both neutralization and recurrence during glioma treatment could induce the alterations of the composition of the gut microbiome.

**Figure 3.**
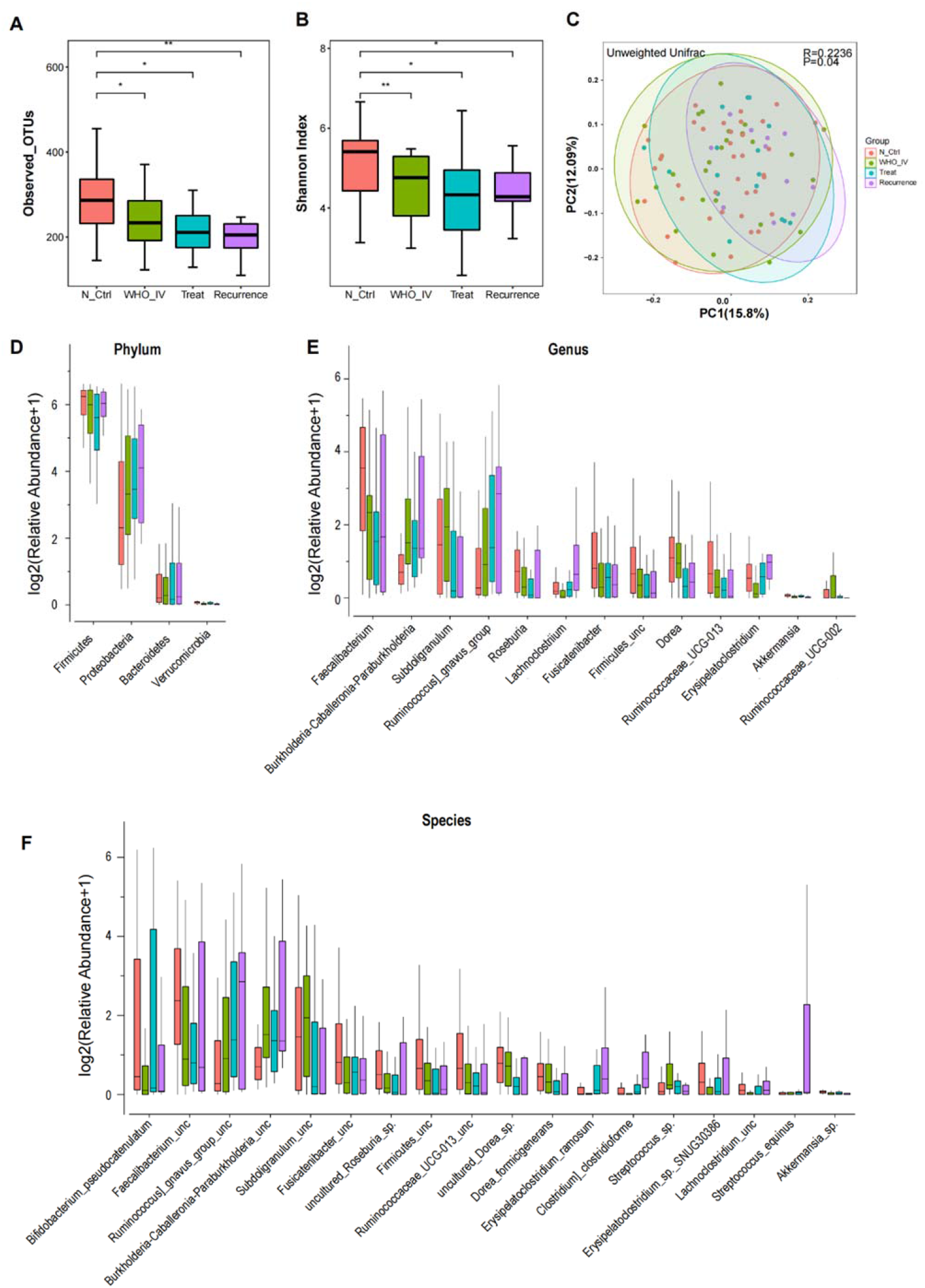
The Alteration of gut microbiota composition in gliomas patients during treatment and recurrence. **(A)(B)** Microbial richness based on number of OTUs_observed, the shannon index. *p<0.05, **p<0.01. **(C)** Principal coordinates analysis (PCoA) for N_Ctrl, WHO_IV, Treat and recurrence groups. Significant differences were observed between patients and controls with ADONIS test. (D) The relative abundance of the main microbiome at the phylum level in different pathological grades groups. (E)(F) The significant changed microbiome in patients with different pathological grades gliomas at the genus and species levels.

We next examined the relative abundances of each phylum, genus and species among these four groups and linear discriminant effect size (LEfSe) was performed. In total, 1 phylum, 13 genera and 18 species were significantly different among these four groups. Despite the low abundance of phyla characteristically present in the gut microbiome, *Verrucomicrobia* which includes many known pathogenic microorganisms differed significantly between these four groups (Figure 3D). It was found to be highly abundant in the primary GBM and Recurrence groups. After removal of tumor mass and continuous adjunctive therapy, relative abundance of several genera from *Firmicutes* decreased. With the recurrence of gliomas, the level of the flora returned to the level of status of the primary tumor. At the genus level (Figure 3E), *Brkholderia-Caballeronia-Paraburkholderia, Ruminococcus-gnavus-gourp* and *Lachnoclostriium* were highly enriched in the Treat and Recurrence groups, whereas *Faecalibacterium, Subdoligranulum, Roseburia*, and an uncharacterized genus of family *Firmicutes* were significantly enriched in N_Ctrl group. At the species level (Figure 3F), species from the *Proteobacteria* and *Verrucomicrobia* phyla were more abundant in the Treat and Recurrence groups. In contrast, species from the *Firmicutes* was significantly decreased in the two groups.

We further performed intergroup analysis among the GBM group, Treat group and Recurrence group. In patients with primary GBM gliomas who received standard therapy, we found that the increased opportunistic pathogens, such as *Streptococcus, Subdoligranulum, Dorea, and Helicobacter*, significantly decreased in the Treat group at the genus level. Probiotics such as *Lachoclostridium* were significantly increased. When the disease progressed, the decreased genera such as *Subdoligranulum* and *Helicobacter*, were further decreased. These results demonstrated that the changes of the gut microbiome were developed during the treatment and recurrence of gliomas.

### Gut metabolism in the fecal metabolome of the primary glioma and healthy control groups

Through above analysis, we demonstrated signature gut microbiome associated with gliomas. We hypothesized that the alterations of the gut microbiome could lead the alterations of metabolites in patients with gliomas. Therefore, we used an untargeted LC-MS-based metabolomics approach to perform metabolome analysis of fecal samples. In total, 5152 detectable peaks were observed in the positive and negative modes defined by retention time and mass/charge. We successfully quantified 2286 metabolites in both the Gliomas and N_Ctrl groups. Principle component analysis (PCA) and partial least squares discriminant analysis (PLS-DA) were applied to observe the differences of distinct groups. As shown in the PCA (Figure 4A) or PLS-DA (Figure 4B), Gliomas and N-Ctrl group were clearly separated into different clusters. Furthermore, we performed the permutation test for PLS-DA (R2=0.7594, Q2=-0.2514, Figure 4C), indicating that the PLS-DA model is reliable without overfitting at the present study. These results indicated broad metabolic shift in glioma carcinogenesis.

**Figure 4.**
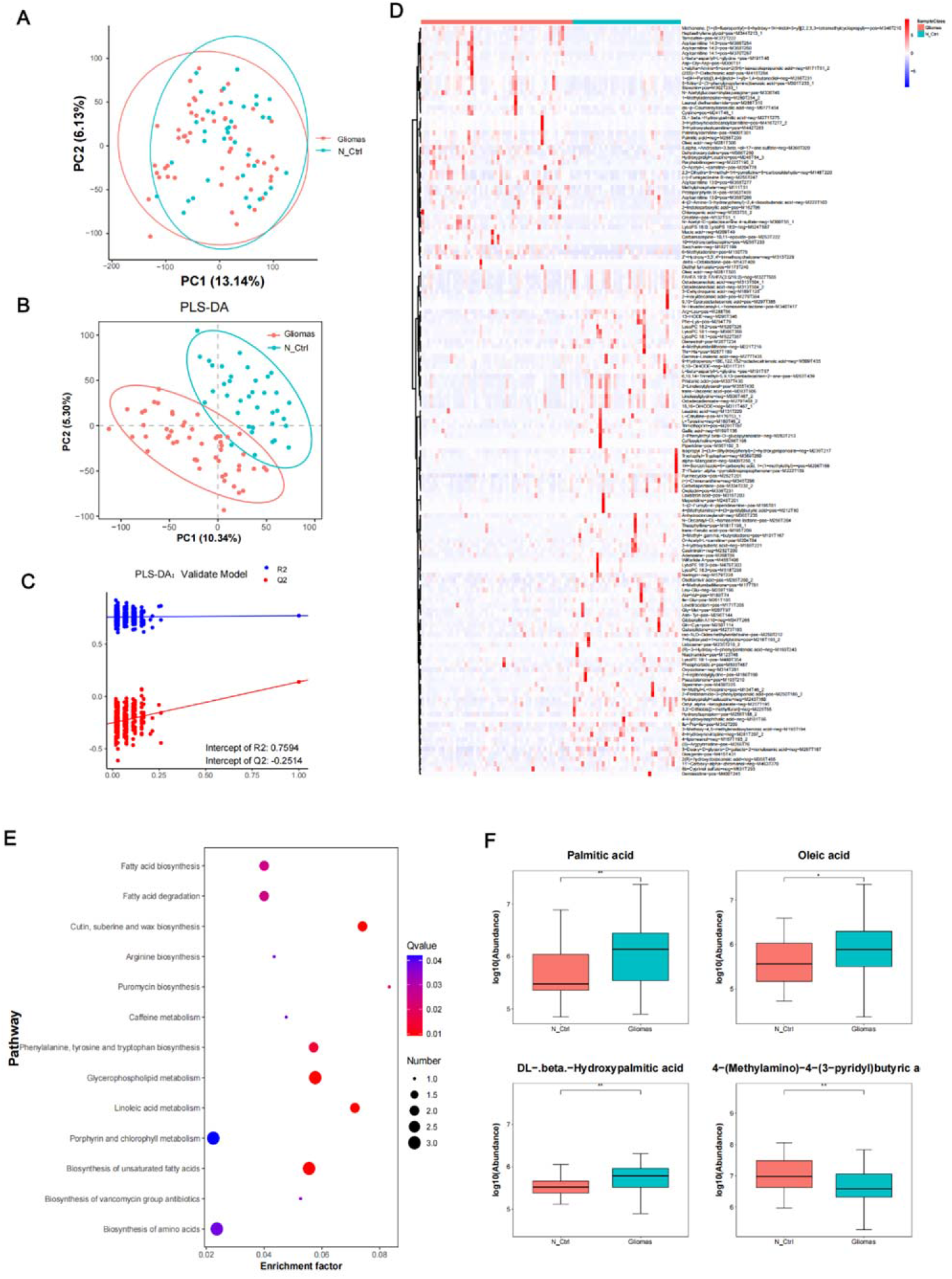
Main discriminatory metabolites identified between Gliomas and N_Ctrl groups. **(A)** PCA score plot. (B) PLS-DA score plot. **(C)** PLS-DA validation plot intercepts. **(D)** The heat map shows the scaled relative intensity of 144 differential metabolites (VIP > 1, p < 0.05). **(E)** Metabolic pathway enrichment analysis of different discriminatory metabolites. (F) Four fatty-acid metabolites were further compared and showing using a box map between Gliomas and N_Ctrl groups. *p<0.05, **p<0.01.

To identify significantly altered metabolites among groups, we analyzed the annotated metabolites of fecal samples. Among the 2286 fecal metabolites, 144 significantly changed metabolites with VIP (variable importance in projection)>0.1, p<0.05, ratio>2 or ratio<0.5 identified in the Gliomas and N-Ctrl group (Table S4), as shown in the Figure 4D. Compared to N_Ctrl group, 50 metabolites were significantly upregulated and 94 metabolites were significantly downregulated in glioma group. The top altered metabolites were 44 lipids and lipid-like molecules, 31 organic acids and derivatives, 15 organo-heterocyclic compounds, 11 organic oxygen compounds, 8 benzenoids and 7 phenylpropanoids and polyketides (Table S4). The majority of carboxylic acids and derivatives and glycerophospholipid metabolites decreased and almost half of fatty acyl metabolites increased. These differential metabolites were mapped onto 43 different KEGG metabolic pathways. The enrichment analysis of the KEGG pathways showed that the most enriched pathways were biosynthesis of unsaturated fatty acids, glycerophospholipid metabolism, linoleic acid metabolism, phenylalanine tyrosine and tryptophan biosynthesis, cutin suberine and wax biosynthesis (Figure 4E). Intriguingly, we found a total of 16 lipids and lipid-like molecules markedly up-regulated in Gliomas samples. KEEG analysis further revealed that unsaturated fatty acids biosynthesis (Figure 4F) was key pathway altered in patients with glioma. We further analyzed the difference abundance of some common fatty acids, such as palmitic acid, with exhibited higher concentrations in gliomas patients than in healthy control groups. In brief, our results suggested dysregulation of lipid metabolisms involved in the development of gliomas.

We further investigated the correlation between the altered abundance of the gut metabolites and the different gut microbiota in gliomas. Spearman’s correlation analysis was used to determine the covariation between 56 OTUs and 144 metabolites. We showed the different metabolites and gut microbiota genera in a heatmap (Figure 5A). We obtained a total of 89 significant associations between differential genera and metabolites. Microbiota and metabolites simultaneously enrich in gliomas or healthy controls presented positive correlations, and vice versa. The correlation networks were applied to illuminate the correlations (r>0.4, Figure 5B). For example, gliomas-enriched genera *Burkholderia-Caballerohia-Paraburkholderia* was observed to be positively associated with several gliomas-specific metabolites such as palmitic acid, acylcarnitine 14:2. The putative mechanism of the positive correlations between genera and metabolites could be explained by microbita producing metabolites. Meanwhile, we found negative association between control enriched genera and gliomas-specific metabolites. Overall, our results indicated a tight interaction of gut microbiota with metabolites in gliomas pathogenesis.

**Figure 5.**
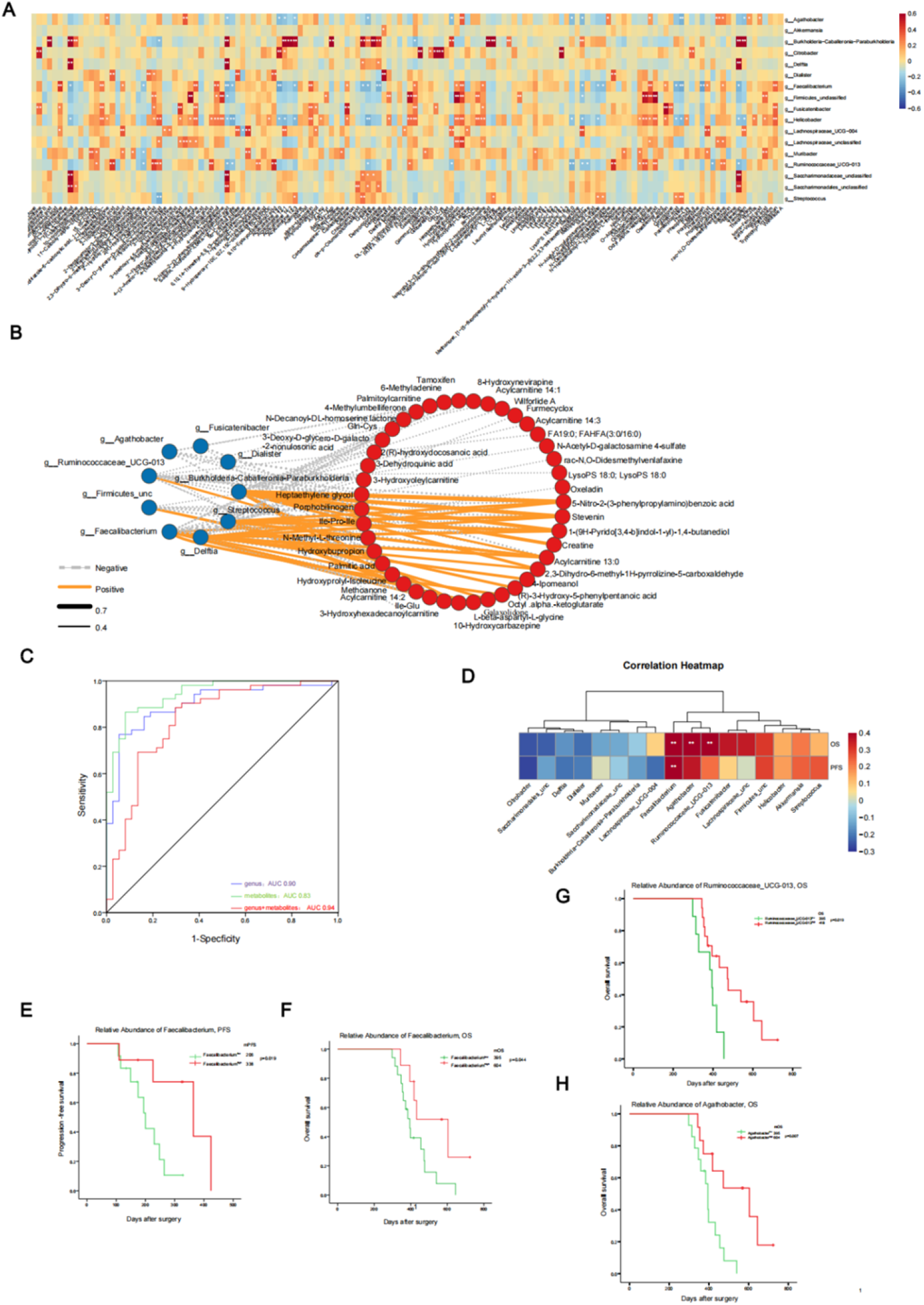
Correlation analysis and potential clinical value of the microbiome and metabolome. (A) Spearman’s correlation analysis was conducted using potential microbiome biomarkers of Gliomas as identified by linear discriminant analysis effect size (LEfSe) and potential metabolite biomarkers identified through multiple linear regression. **(B)** The correlation among differential metabolites and differential genera is shown using a cooccurrence network. In the network, Spearman correlation coefficient values below −0.4 (negative correlation) are indicated as gray edges, and coefficient values above 0.4 (positive correlation) are indicated as red edges. (C) Predictive models based on microbiota and metabolites for gliomas. (D) Spearman’s correlation analysis was calculated to determine correlations between fecal bacterial genera and PFS or OS. Hierarchical clustering was performed with bacterial genera that included the correlation coefficient with p-value <0.05. *p < 0.05; **p < 0.01. (E) The subjects were divided into groups according to abundance of intestinal *Faecaliacterium* Plots of PFS (**E**) and OS (F) were drawn by the Kaplan– Meier method. (G) The subjects were divided into groups according to abundance of intestinal *Ruminococcaceae UCG_13* Plots of OS. (H) The subjects were divided into groups according to abundance of intestinal *Agathobacter* Plots of OS.

### Gut microbiome and metabolites identified and associated with outcomes of glioma patients

We investigated the potential value of the gut microbiota combined with metabolites for identifying gliomas from healthy controls, we used the random forest (RF) algorithm to construct a promising predictive model. For identifying gliomas from healthy controls, 8 gut microbiota genera (*Helicobacter, Burkholderia-Caballeronia-Paraburkholderia, Akkermansia, Fusicatenibacter, Faecalibacterium, Delftia, Streptococcus* and R*uminococcaceae_UCG-013*) were selected with an area under the curve (AUC) of 0.901 (Table S5, Figure 5C). Among these genera, *Helicobater, Akkermansia, Streptococcus, Burkholderia-Caballeronia-Paraburkholderia* were more enriched in glioma patients, *Delftia, Fusicatenibacter, Faecalibacterium, Ruminococcaceae_UCG-013* were enriched in the gut of healthy controls. We obtained 10 metabolites (methyladenine, saccharin, 3-methoxy-methylenedioxybenzoicacid, theophylline, linoleoylglycine, transferulicacid, palmitic acid, linoleoylglycerol, 13-HODE and octadecaedioic acid) as markers to distinguish gliomas from healthy controls with an area under the curve (AUC) of 0.832 (Table S5, Figure 5C). To get better discrimination, we combined these 8 different genera and 10 metabolites to detect gliomas. More accurately result was obtained by an AUC of up to 0.944. These results indicated that the gut microbiome and metabolites could be valid predictors for gliomas.

Through sequencing of 16S rRNA amplicons, it is reasonable that the altered gut microbiome and metabolites could be associated with prognosis of patients with gliomas. We investigated inter-associations between the microbial genera of Glioblastoma multiforme (GBM, WHO IV) and PFS or OS by spearman’s correlation analysis. At the genera level, relative abundance of *Faecalibacterium* in the gut microbiota was positively correlated with both PFS (r=0.60, p=0.002, Figure S3A) and OS (r=0.52, p=0.008,Figure S3B). OS also was positively associated with *Agathobacter* (r=0.52, p=0.006,Figure S3D) and *Ruminococcaceae-UCG-013* (r=0.50, p=0.010, Figure S3C) (Figure 5D). Even though several generas were negatively associated with PFS or OS, there were no statistical significance. We stratified the GBM patients into a high abundance group and a low abundance group based on these enriched taxa in WHO_IV group. Survival analysis showed that patients with high proportions of fecal *Faecalibacterium* before therapy had significantly better PFS than those with low proportions of *Facecalibacterium* (mPFS: *Facecalibacterium*^high^ 320 days, *Facecalibacterium*^low^ 281 days, p=0.005, Figure 5E). OS also was longer in patients with fecal *Facecalibacterium*^high^ than in those with *Facecalibacterium*^low^ (mOS: *Facecalibacterium*^high^ 604 days, *Facecalibacterium*^low^ 395 days, p=0.044, Figure 5F). In multivariate analysis, *Facecalibacterium* was the significantly predictive factor associated with PFS, with hazard ratios of 4.35 (95%CI: 1.42-13.28, p=0.010), but no statistical significance with OS (p=0.053). *Agathobacter* and *Ruminococcaceae_UCG-013* enrichment was also associated with better OS. GBM patients enriched with high *Agathobacter* had significantly better OS (mOS: 604 vs 395 days, p=0.007, Figure S5H). Moreover, patients with GBM harboring a high abundance of *Ruminococcaceae_UCG-013* had significantly prolonged OS (mOS: 418 vs 395 days, p=0.019, Figure 5G). The fecal metabolites were greatly influenced by the environment and dietary habits, making it challenging to assess the correlations of metabolites with PFS or OS.

### Gliomas lead to alterations in the gut microbiome in a glioma mouse model

To determine whether glioma leads to dysbiosis of the gut microbiome, we conducted the experiment in two groups, the sham-operation (control) group and the GL261 cell implantation (tumor) group (Figure 6A). We collected fecal samples before glioma implantation or 3 weeks after glioma implantation, and 16S rRNA gene sequencing analysis was used to evaluate the impact of glioma on the gut microbiome. We compared the richness and diversity of the gut microbiome between the two groups (control and tumor groups). We found significant differences in the Observed_otus (p=0.023) (Figure 6B), Shannon index (p=0.032) (Figure 6C), Chao1 (p=0.028) and Simpson index (p=0.015) (Figure S2A and B). PCoA analysis for beta-diversity did not show any significant difference between the two groups (p=0.15, Figure 6D). Similar to humans, the prominent phyla were *Firmicutes, Bacteroidetes, Actinobacteria* and *Proteobacteria* (Figure 6E). We also compared the F/B ratio. In glioma models, the ratio changed, which indicates dysbiosis following tumor development, but the change was not statistically significant (Figure S2C).

**Figure 6.**
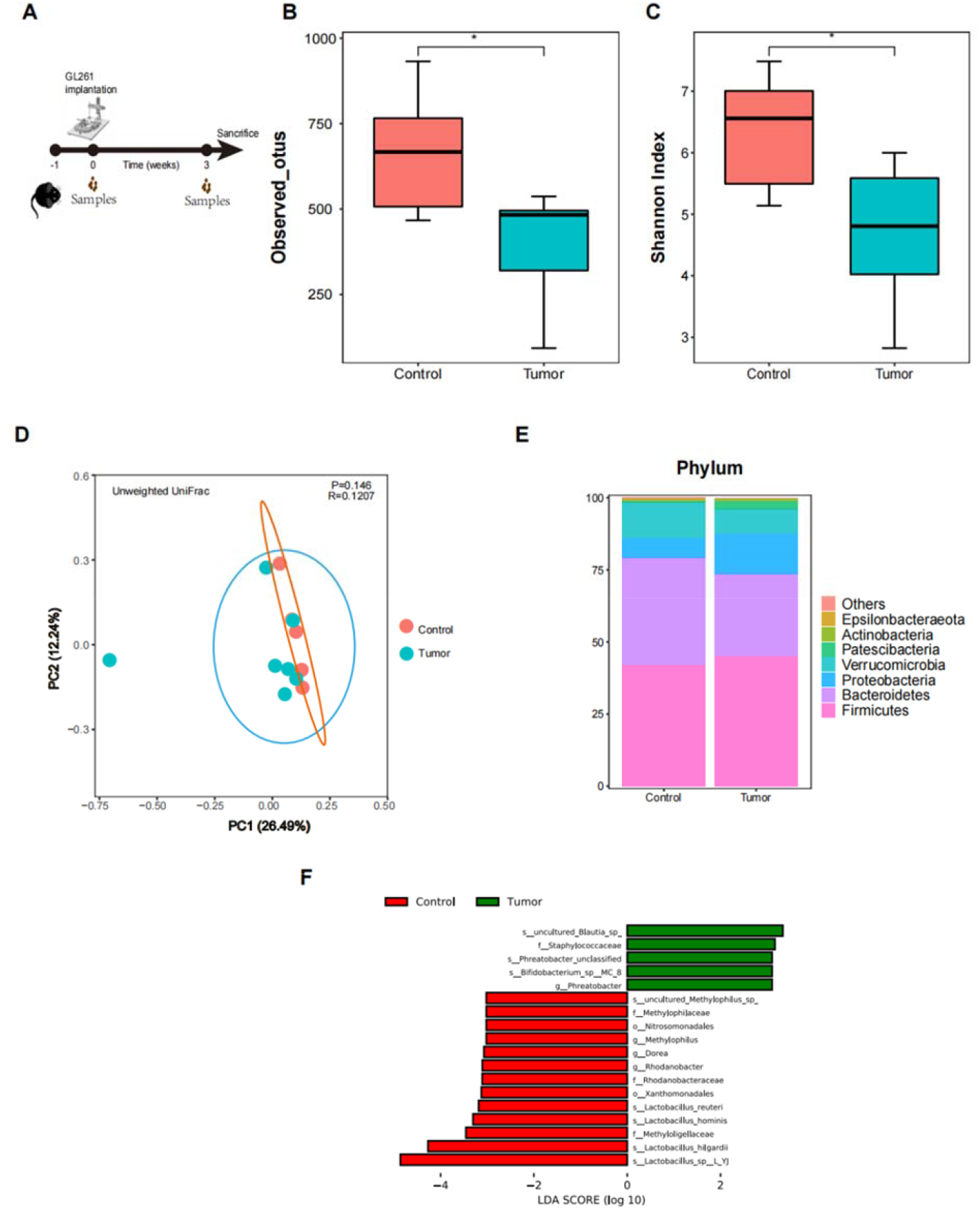
Alterations in gut microbiota composition in glioma models. (A) Schematic diagram showing the experimental design and timeline of the female C57BL6/N mouse glioma model. (B) and (C) Microbial richness based on the number of OTUs observed and the Shannon index. *p<0.05. (D) Principal coordinates analysis (PCoA) for the control and tumor groups. Nonsignificant differences were observed between patients and controls with the ADONIS test. The first two principal coordinates (PCs) were each labeled with the percentage of variance explained. (E) The relative abundance of bacterial phyla clustered into different groups was determined. (F) LDA scores for the bacterial taxa differentially abundant between the control and tumor (glioma models) groups (LDA > 3.0), generated by LEfSe analysis. Red bars indicate taxa that were enriched in the control group, and green bars indicate taxa that were enriched in the tumor group.

Then, we compared the compositions of the gut microbiota in the tumor group with those in the control group to identify the specific communities related to glioma by using LEfSe analysis (Figure 6F). In total, LEfSe analysis revealed 18 discriminative OTUs (LDA>3.0, p<0.05, Figure 6F). In glioma models, the gut microbiome was dominated by *Blautia, Phreatobacter, Bifidobacterium* and *Phreatobacter*, whereas in the control group, *Lactobacillus, Rhodanobacter, Dorea* and *Methylophilus* were enriched at the genus level. Amazingly, these discriminative microbiotas were also found to be enriched in the human gut microbiome of glioma patients. Therefore, the results showed that the alterations of the gut microbiome accompanied by gliomas in the two different species were similar.

### Fecal microbiota transplantation (FMT) from glioma patients with high levels of *Faecalibacterium* protects against tumor growth

Through clinical analysis, we demonstrated that glioma patients with high levels of *Faecalibacterium* could obtain better prognoses. Thus, we performed fecal microbiota transplantation to validate this result. We chose fecal samples from two glioma patients with good prognosis and high levels of *Faecalibacterium*, of which the relative abundance was 32% and 24% (as the H FMT group). Fecal samples from two patients with lower survival time and low level of *Faecalibacterium* (the relative abundance was 0.1% and 0.5%) (as L FMT group). All these patients were marched by age and sex. To avoid the impact of the endogenous microbiota, we pretreated mice with ABX for 7 days before FMT (Figure 7A). Three weeks after implantation, MRI was conducted. H FMT restricted tumor growth compared to the L FMT group (Figure 7B and C). Notably, H FMT also had a significant impact on survival (Figure 7D). Compared to L FMT, H FMT induced a decrease in tumor cell proliferation as measured by Ki-67 staining (Figure 7E and F). Neither treatment altered the percentage of apoptotic (as measured by caspase 3 staining) tumor cells (Figure 7E and G). Overall, these data showed that a high level of *Faecalibacterium* was able to protect against tumor growth.

**Figure 7.**
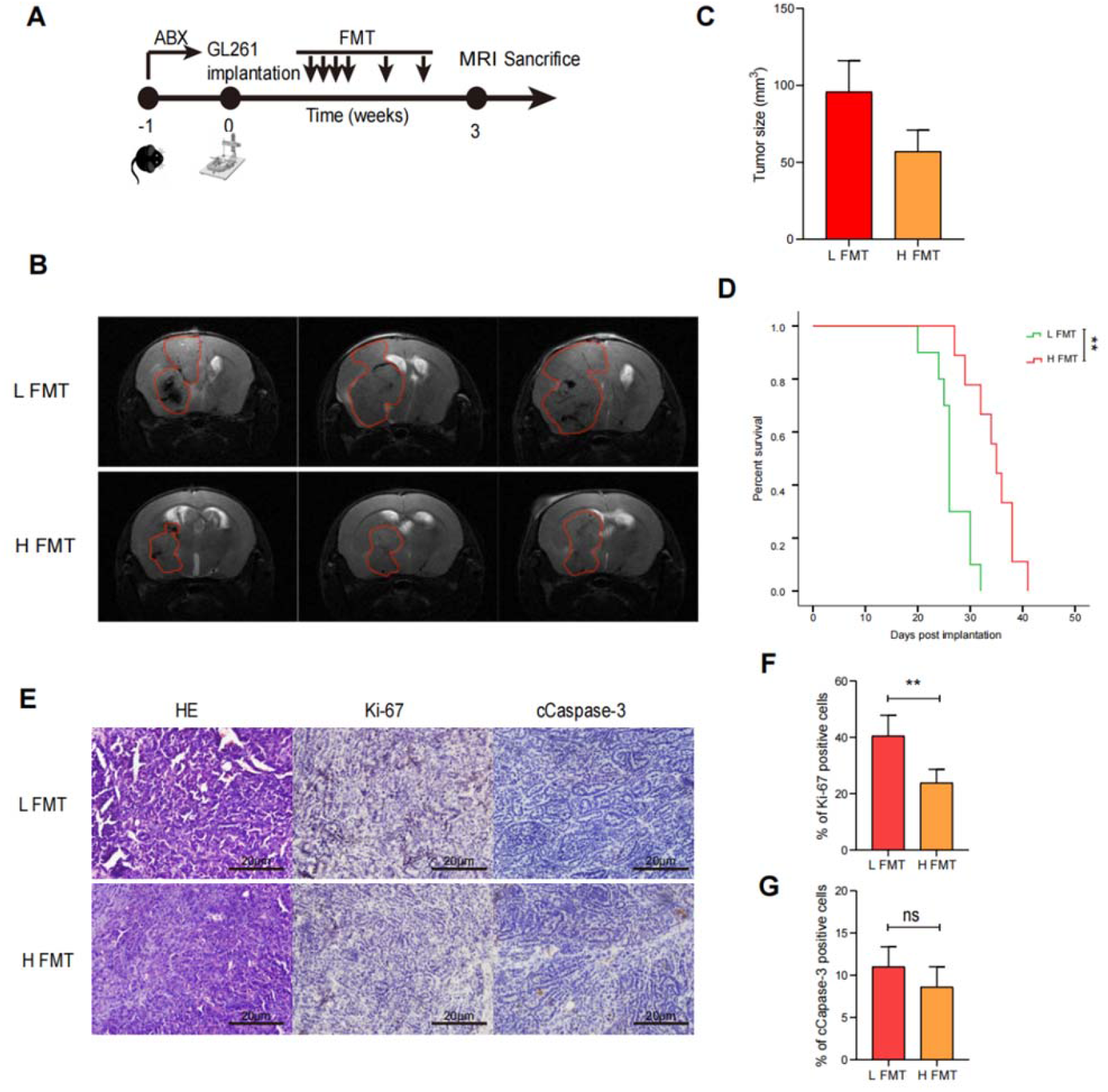
Fecal microbiota transplantation from glioma patients impacts tumor growth in glioma model mice. (A) Schematic diagram showing the experimental design and timeline of the female C57BL6/N mouse glioma model. (B) Experimental scheme and representative MRIs of L FMT and S FMT mice at 3 weeks after GL261 cell implantation. (C) Tumor size of different groups. (D) Survival curve. **p<0.01. (E) Representative hematoxylin and eosin (H&E), Ki-67, and Caspase-3 immunohistochemical staining. Scale bar, 20 μm. (F) and (G) Quantification of Ki-67-positive cells (F) and caspase-3-positive cells (G). **p<0.01.

## Discussion

Many factors, including genetic and environmental factors, can be the causes of gliomas. Nowadays, more and more evidences indicate that the gut microbiome which is an important part of the host environment, is closely related to diseases of the CNS, such as Parkinson’s disease(19) and mental disorders(18, 27). Gliomas are the most common diseases in CNS. We conducted a pilot exploration to characterize the gut microbiome and metabolites in fecal samples from patients with gliomas. We demonstrated the profile changes of the gut microbiome and its metabolites in glioma patients and identified those microbiota and metabolite candidates that might contribute to the development of gliomas. In addition, the identified microbial biomarkers exhibited remarkable power to differentiate gliomas from healthy people and were associated with the prognosis of gliomas. We also found the alterations of gut microbiome structure in mice with gliomas. Moreover, we performed FMT experiments to verify the prognosis value of *Faecalibacterium*. All these findings provide further understanding of the gut-brain axis.

In our study, we observed lower bacterial community richness (Chao1 index) and no significant difference in community diversity (Simpson index) in patients with gliomas compared with healthy controls. Similar results were found in patients with gliomas of different stages. The unweighted Unifrac index of the Gliomas group was statistically lower than that of the healthy control group. Several previous studies suggested that the decreased microbial diversity of the gut seems to be associated with unhealthy conditions. These findings are consistent in extraintestinal diseases such as breast cancer(15, 28) and liver cancer(14). The animal study by Anthony Patrizz et al(24) suggested that the diversity of the gut microbiome was lower in gliomas mice models than in healthy controls. Different pathological stages of thyroid neoplasm also showed that the diversity of the gut microbiome significantly changes(29). Moreover, postoperative therapy influenced the composition of the gut microbiome as well. In a study of non-small cell lung cancer patients receiving PD-1 (anti-programmed death 1) exhibited higher gut microbiome diversity(11, 30). Similar to other related studies(24), our subsequent animal study also found that the gut microbiota was disordered in glioma-bearing mice. We propose that this change in gut microbiota is related to the health status of the host. Both glioma patients and tumor-bearing mice have the possibility of multiple organ dysfunction during their survival compared with healthy controls. The “second genome” gut microbiota is affected by the growth of glioma. Overall, the differences in alpha diversity and beta diversity suggested that changes of the gut microbiome in glioma patients, which are associated with the burden of the tumor and tumor grades.

As our study showed, *Firmicutes, Bacteroidetes, Actinobacteria*, and *Proteobacteria* were the predominant phyla both in glioma patients and healthy adults, also in model mice. The F/B (Firmicutes/ Bacteroidetes) ratio is considered representative of the health status and may reflect the ecological equilibrium of the gastrointestinal tract. Some studies have showed that the F/B ratio increases or decreases in diseases(31, 32). Anthony Patrizz et al(24) also found that the F/B ratio was lower in glioma mice than that in healthy controls. In this present study, we found a change of the ratio, but with no significant difference. However, compared to the healthy controls, we confirmed that the structure of the gut microbiome was altered with the development of gliomas. Several opportunistic pathogens were significantly overrepresented (all LDA scores >3.0) in the glioma group. One of them is the *Streptococcus*. Many studies have shown that they are also enriched in some other tumors such as colorectal carcinogenesis(33), and lung cancer(34). *Streptococcus* is associated with carcinogenesis by enhancing the expression of proliferation markers. In addition, *Faecalibacterium* and *Ruminococcaceae* as probiotics were decreased in glioma patients in the current study. These genera have been identified to produce short-chain fatty acids (SCFAs)(35), such as acetylic acid, propionate and butyric acid. To the best of our knowledge, SCFAs have direct or indirect anti-tumor effects(25, 36). Furthermore, we also found that probiotics were decreased in glioma-bearing mice. A variety of *Lactobacillus* (such as *Lactobacillus reuteri* and *Lactobacillus hominis*) with anti-inflammatory effects were reduced in glioma-bearing mice, which could affect the growth of a variety of tumors(37, 38).Moreover,we found that *Akkermansia* which is a notable microorganism, was increased in patients with gliomas. Several studies have reported an increased abundance of *Akkermansia* in patients with CNS disorders, such as Parkinson’s disease, and multiple sclerosis(39, 40). These results indicate that chronic inflammatory changes in the CNS are accompanied by the growth of *Akkermansia*. In our subgroup analysis, we found some microbiomes associated with IDH-1 and MGMT status. The mechanisms of these associations remain unclear. We hypothesized that the possible mechanism could be generated through participation in lipid metabolism and anti-inflammatory metabolites. Certainly, further cell and animal experiments are needed to confirm the causal relationship.

Of note, we investigated the profile of gut microbiome changes from detection, treatment to recurrence of gliomas. In the treatment and recurrence of gliomas, species from the *Proteobacteria* and *Verucomicrobia* were enriched. Compared to GBM, several opportunistic pathogens, such as *Streptococcus* and *Helicobacter* significantly increased and several probiotics such as *Lachoclostridium* were increased. Disputed with the results from Anthony Patrizz et al(24), the structure of gut microbiome did not recover normally. Similar to GBM, members from *Proteobacteria* more enriched in recurrence of gliomas. Future efforts with more samples are necessary to investigate the changes of gut microbiome during the treatment and recurrence of gliomas.

As mentioned above, we discovered that the altered gut microbiota genera in the glioma group were significantly correlated with lipid-related metabolites. In our study, the fecal metabolic analysis identified 144 altered metabolites, mainly including lipid and lipid-like metabolic molecules, some amino acids, flavonoids, benzoic acids and derivatives. The interruption of lipid metabolism is related to the occurrence and development of some types of human cancers, including colorectal carcinoma and breast cancer. Previous studies investigating glioma metabolomics mainly involved tissue specimens, serum and CSF samples. One study investigated lipidosis signatures in ectopic and orthotopic human GBM xenograft models and identified glycosphingolipids, glycerophosphoethanolamines triacylglycerols, and glycerophosphoserines as four main classes of lipids with greatest fold effect(41). Another study that investigated serum metabolism in glioma noted LDL, unsaturated lipids, cholesterol and other sphingolipid metabolites(42). Some of these metabolites were also identified in our analysis. Therefore, we speculate that the gut microbiota dysbiosis is closely related to the development of glioma, and that lipid metabolism dysregulation may be a key factor. In reality, cancer cells are specifically altered in several aspects of lipid metabolism to meet their high bioenergy requirements(43). By remodeling lipid metabolism, tumor cells can enhance their aggressiveness and enhance their role in cell growth, proliferation and other mechanisms. Lipids are abundant in the cerebral microenvironment. They not only maintain the normal function of astrocytes, but also plays a certain role in the progression of glioma, and this role is closely related to the overall lipid metabolism of the body. In addition, consistent with microbial changes, we also found that the levels of SCFAs were decreased in gliomas groups.

Beyond the simple description alterations of gut microbiome in patients with gliomas, our study proposes the idea of gut microbiome and metabolites-based predictors for gliomas by multivariate analysis. We find that this predictor is more accurate. Several of 8 gut microbiota genera and 10 metabolites markers have been reported to functionally impact upon the host. *Helicobacter* and *Streptococcus* can affect immune maintenance and pro-infammation through recruiting and regulating differentiation of immune cells in intestinal(44). Methyladenine can be involved in RNA and DNA synthesis. Emerging evidence shows that the gut microbiome can modulate the efficacy of pharmaceutical agents in the therapy of tumors(45, 46). We also found significantly positive and negative predictive correlations with the PFS and OS of patients with gliomas. Patients with high abundance of *Faecalibacterium, Ruminococcaceae_UCG-013* and *Agathobacter* had a better prognosis than patients with a lower abundance. Members of these three genera are probiotics, and achieve anti-cancer effects by producing secondary metabolites and reducing the inflammation. Through analysis we found the reduced abundance of *Faecalibacterium, Ruminococcaceae_UCG-013*, and *Agathobacter* in GBM patients, compared to healthy controls, but only the alteration of *Ruminococcaceae_UCG-013* had statistical significance. The main metabolites of these three genera are SCFAs, inhibitors of HDACs, which could repress the development of gliomas. *Faecalibacterium* also can exert anti-inflammatory functions by inhibiting IL-8 secretion(47). Nevertheless, the overall role of the gut microbial community is thought to have an effect on the survival period of patients. To test whether such an effect exists, we fed human stool to intracranial tumor-injected mice to study the effect of gut microbiota on glioma. Stool from patients with shorter survival periods promoted tumor growth and shortened survival periods in tumor-bearing mice. This suggested that the stool content was an important factor in determining the phenotypes. In fact, microbial communities such as CRC can directly contact tumors and promote their growth through immune and inflammatory pathways, and there is a causal relationship between them(48).As a distant organ, the central nervous system and the gut may communicate through the “gut-brain axis”, which requires metabolites and neurotransmitters as messengers(8, 26). According to our study, patients with shorter survival periods had more severe dysbiosis, while other similar studies showed that dysbiosis showed a decrease in adaptive immunity, a decrease in immune killer cells in the tumor microenvironment, and an increase in proinflammatory cells and cytokines(47, 49). These changes were also accompanied by changes in metabolite levels, such as a decrease in SCFAs and an increase in LCAs(47) in late-stage glioma. The survival time of patients is related to many factors, and the mechanism is complex, but we confirmed that there is a causal relationship between intestinal microbiota and glioma growth and patient survival time through experiments.

It is noteworthy to mention that our study provides a novel insight into the potential contribution of the gut microbiome and metabolites to gliomas. However, there were several limitations in this study. First, this study was only a single-center cross-sectional study, so the sample size was limited. Second, the composition of the gut microbiome and metabolites could be affected by several confounding factors varying from different individuals, such as geographical conditions, and dietary habits. Third, in this study we used 16S rRNA gene sequencing to investigate the composition of the gut microbiome. However, it could be insufficient to illustrate the whole profile of the gut microbiome. Deep gut microbiome profiling techniques such as shotgun sequencing of metagenomics and meta transcriptomics are available. To investigate the metabolites of the fecal samples, nontarget metabolomics was used, but detailed information on specific metabolic pathways could not be found. Therefore, further studies with large samples using metagenomics sequencing and target metabolomics are needed to investigate the potential causal mechanisms of the association between the composition of the gut microbiome, fecal metabolites and gliomas.

In conclusion, this study revealed that the development of gliomas could make the alterations of the gut microbiome at the phylum, genus and species levels and metabolites. The presence of a specific gut microbiome and metabolites may act as biomarkers to discriminate gliomas. We also found that specific gut microbiota may contributes to prognosis. These findings provide insights into the relationship between the gut microbiome, metabolites and gliomas. This finding paves the way for further research in clinical practice.

## Materials and methods

### Ethics statement

This study was reviewed and approved by the Institutional Review Board at Nanfang Hospital, Department of Neurosurgery, Nanfang Hospital of Southern Medical University (Guangzhou, China). All animal procedures were performed in accordance with the NIH Guidelines for the Care and Use of Laboratory Animals (Bethesda, MD).

### Study design and fecal sample collection

A total of 51 patients diagnosed with primary gliomas (Gliomas group), 15 patients who underwent TMZ chemoradiotherapy (Stupp regimen) (Treat group), 11 patients with recurrent gliomas after chemoradiotherapy (Recurrence group) and 37 healthy volunteers for conventional medical examination (N_Ctrl group) from Nanfang Hospital, Southern Medical University (Guangzhou, China) from January 2020 to December 2021 were recruited for this study. All participants were Han Chinese and from Guangdong Province. The research was performed in accordance with the principles of the Declaration of Helsinki and Rules of Good Clinical Practice. Written informed consent was obtained from all the patients and healthy volunteers. All the enrolled patients underwent craniotomy and tumor resection, and glioma was confirmed by pathological examination according to the 2016 World Health Organization classification of tumors of the CNS. For healthy volunteers, physical examinations, such as routine examination of blood, feces, urine, renal and liver function, were used to identify any serious diseases. None of the participants in this study had a history of gastrointestinal disorders, autoimmune diseases, diabetes, malignant tumors or other severe diseases. Individuals who received antibiotics, probiotics or prebiotics in the previous 3 months were excluded.

Comprehensive clinical information for the enrolled patients and healthy volunteers, including age, sex, body mass index (BMI), tumor grade, tumor pathological type and smoking status was collected. Fecal samples from the study subjects were collected in sterilized 7 ml tubes, transported to the laboratory and frozen at -80°C immediately until DNA extraction. This study was approved by the Institutional Review Board at the Nanfang Hospital of Southern Medical University.

We collected the follow-up information of the WHO IV glioma patients from Gliomas group. Progression-free survival (PFS) was defined as the time from diagnosis until recurrence, death, or the last follow-up date if the patient showed no disease progression or death. Overall survival (OS) was defined as the time from diagnosis until death or the last follow-up date in which the patient did not die. Patients who were alive at the date of the final analysis were censored at the last follow-up date. The updated Response Assessment in Neuro-Oncology (RANO) criteria was used to define tumor progression or recurrence(44).

### DNA extraction and 16S rRNA gene sequencing

DNA from different samples was extracted using the E.Z.N.A ®Stool DNA Kit (D4015, Omega, Inc., USA) according to the manufacturer’s guidelines. Two percent agarose gel electrophoresis was used to verify DNA integrity and size. AmPure XT beads (Beckman Coulter Genomics, Danvers, MA, USA) were used for DNA purification and Qubit (Invitrogen, USA) was used for DNA qualification. The V3-V4 hypervariable region of the prokaryotic (bacterial and archaeal) small-subunit (16S) rRNA gene was amplified with primers 341F (5’-CCTACGGGNGGCWGCAG-3’) and 805R (5’-GACTACHVGGGTATCTAATCC-3’)(50). The amplicon pools were prepared for sequencing and the size and quantity of the amplicon library were assessed on an Agilent 2100 Bioanalyzer (Agilent, USA) and with the Library Quantification Kit for Illumina (Kapa Biosciences, Woburn, MA, USA), respectively. The libraries were sequenced on a NovaSeq PE250 platform.

### 16S rRNA gene sequencing data analysis

The samples were sequenced on an Illumina NovaSeq platform according to the manufacturer’s recommendations, provided by LC-Bio. Paired-end reads were assigned to samples based on their unique barcode, truncated by cutting off the barcode and primer sequence, and were merged using fast length adjustment of short reads (FLASH). Quality filtering of the raw reads was performed under specific filtering conditions to obtain the high-quality clean tags according to the fqtrim (v0.94). Chimeric sequences were filtered using Vsearch software (v2.3.4). After dereplication using DADA2, we obtained a feature table and feature sequence. The sequence alignment of species annotation was performed by using Blast, and the alignment database were SILVA and NT-16S. Alpha diversity and beta diversity were calculated by QIIME2, in which the same number of sequences was extracted randomly by reducing the number of sequences to the minimum of some samples, and the relative abundance was used in bacterial taxonomy. Alpha diversity was measured to identify the complexity of the species diversity of every sample. Principal coordinates analysis (PCoA) was used to estimate beta diversity. The Wilcoxon rank-sum test was used to compare bacterial abundance and diversity. Linear discriminant analysis effect size (LEfSe) was performed to evaluate the differentially abundant taxa.

### Extraction of fecal metabolites and LC-MS analysis

The fecal samples were thawed on ice, and metabolites were extracted with 50% methanol buffer. Briefly, 20 μL of sample was extracted with 120 μL of precooled 50% methanol, vortexed for 1 min, and incubated at room temperature for 10 min; the extraction mixture was then stored overnight at -20°C. After centrifugation at 4,000 g for 20 min, the supernatants were transferred into new 96-well plates. The samples were stored at -80°C prior to the LC-MS analysis. In addition, pooled QC samples were also prepared by combining 10 μL of each extraction mixture(51).

All the samples were acquired by the LC-MS system following machine orders. All chromatographic separations were performed using a Thermo Scientific UltiMate 3000 HPLC. An ACQUITY UPLC BEH C18 column (100 mm*2.1 mm, 1.8 μm, Waters, UK) was used for the reversed phase separation. Metabolites eluted from the column were detected by a high-resolution tandem mass spectrometer Q-Exactive (Thermo Scientific).

The LC-MS raw data files were converted into mzXML format and then processed by XCMS, CAMERA and metaX software (). Each ion was identified by combining retention time (RT) and m/z data. The online KEGG, and HMDB databases were used to annotate the metabolites by matching the exact molecular mass data (m/z) of the samples. We also used an in-house fragment spectrum library of metabolites to validate the metabolite identification. PCA was performed for outlier detection and batch effect evaluation to interpret cluster separation. Supervised partial least squares discriminant analysis (PLS-DA) was conducted through metaX to discriminate the different variables between groups. The pathway analysis of different metabolites was conducted with MetaboAnalyst.

### Animal experiments

#### Cell line and glioma models

The protocols for the cultivation of the mouse GBM cell line (GL261) and glioma models are provided in the **Supplemental Materials and Methods** (available online).

#### MRI experiments

Intracranial tumor growth was monitored in vivo in isoflurane-anesthetized mice by MRI after inoculation using a Bruker 7.0 T scanner (Bruker BioSpin GmbH). T2-weighted images were acquired by a rapid acquisition relaxation-enhanced sequence.More details are provided in the **Supplemental Materials and Methods** (available online).

#### Histology/histopathology

Tumor samples from mice were embedded in paraffin as routine methods, and tissue section staining was performed as previously described. Anti-Ki67 and anti-Cleaved Capase3 were used. Two different investigators reviewed and scored the stained tissue sections separately.the details of the staining and the scoring system for determining the percentage of positive cells and staining intensity are available in the **Supplemental Materials and Methods** (available online).

#### FMT experiments

To eliminate the effects of the endogenous intestinal microbiome, mice were treated with ABX (a cocktail of vancomycin 0.5 g/L, ampicillin 1 g/L, streptomycin 1 g/L and metronidazole 1 g/L) in the sterile drinking water of mice. The solution was changed every two to three days, and the bottles were changed once a week. Before fecal microbiota transplantation (FMT), mice were treated with ABX for 7 days, and then GL261 cells were implanted. After transplantation, thawing fecal materials from the selected glioma patients or healthy humans were applied to each mouse. A total of 200 μl of the suspension was transferred into each ABX-pretreated mouse by oral gavage. Mice received FMT for four consecutive days on the first week and once a week for the following weeks.

### Statistical analysis

Statistical analyses were performed using SPSS 19.0 Software (San Diego, CA), QIIME and R software (v3.5.2). For normally distributed data, the data are expressed as the mean ± SD and the median for nonnormally distributed data. T tests and Wilcoxon rank-sum tests were used to evaluate the differences between groups. We used the Spearman’s rank correlation analysis to examine the correlation between clinical indexes and species or between species and metabolites. The random forest algorithm was used to produce a model of the microbiota and metabolites based on the mean decrease accuracy (MDA). Through this method, some different microbiota and metabolites were selected as the potential features for the predictive model. Receiver operating characteristic (ROC) curves were utilized to evaluate the performance of the features. Spearman’s rank correlation test was conducted to explore the association between the gut microbiota and PFS or OS. Cox proportional hazards regression analysis and Kaplan-Meier analysis were used to calculate the cumulative survival proportions. A p value <0.05 was considered statistically significant.

### Data Availability Statement

Some or all data, models, or code generated or used during the study are available in a repository or online in accordance with funder data retention policies. (DOI:10.5281/zenodo.6650755.URL:https://doi.org/10.5281/zenodo.6650755)

## Author contributions

Y.T.L., S.T.Q., M.Z., C.S., L.Y.S., and J.W.G designed the study, M.Z., C.S., J.J.L, T.W. and H.L. conducted the study, enrolment and managed the patients. M.Z., C.S. and L.Y.S. collected the clinical data and samples. M.Z, C.S., T.W. and L.W.Z performed the 16rDNA sequencing, LC-MS experiments and bioinformatic study. M.Z., T.W. and Y.T.L. performed the data analysis. Y.T.L., S.T.Q., M.Z., C.S., L.Y.S. and H.L. discussed the data. M.Z. and C.S. wrote the manuscript.

---

**Figure S1.**
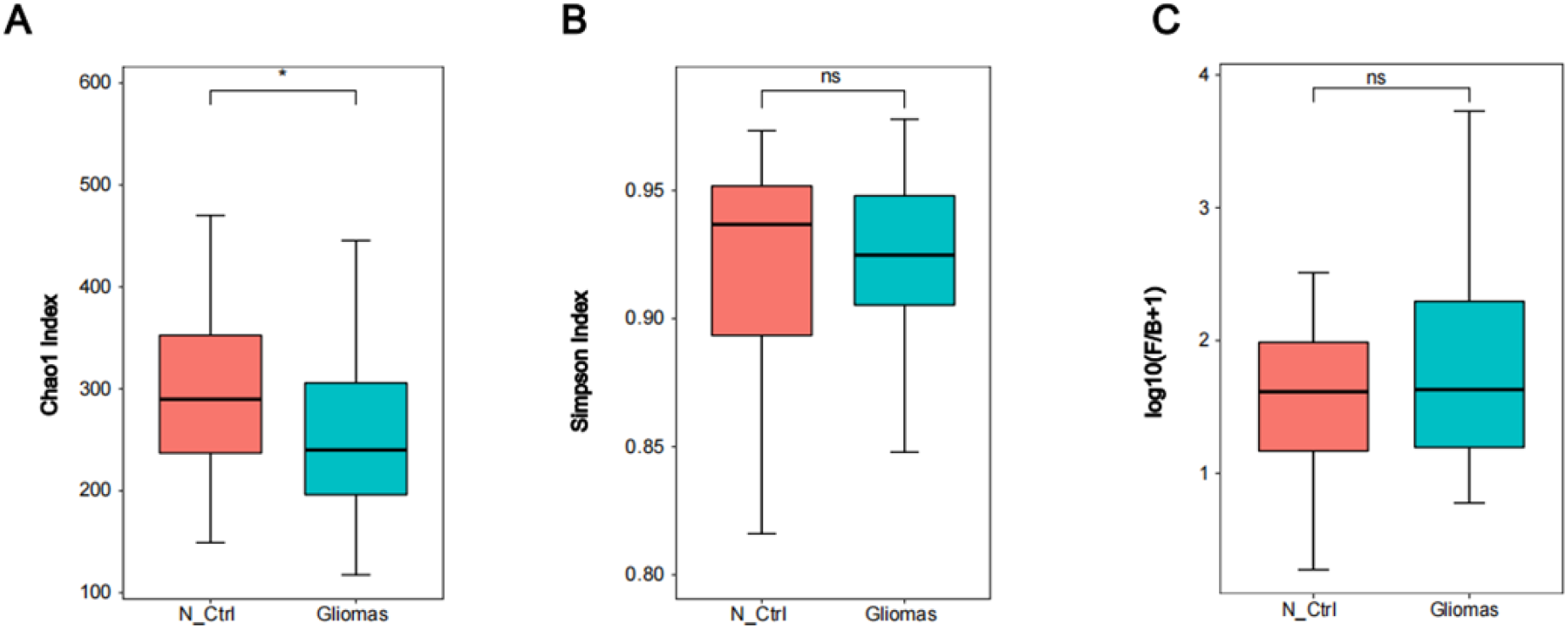
Relative bacterial richness analysis between Gliomas and N_Ctrl groups. **(A)(B)** Microbial richness based on number of the chao1 index, the simpson index. *p<0.05, ns: no significant. (C) The *Firmicutes*/*Bacteroidetes* ratio showed no significant variation between Gliomas and N_Ctrl groups.

**Figure S2.**
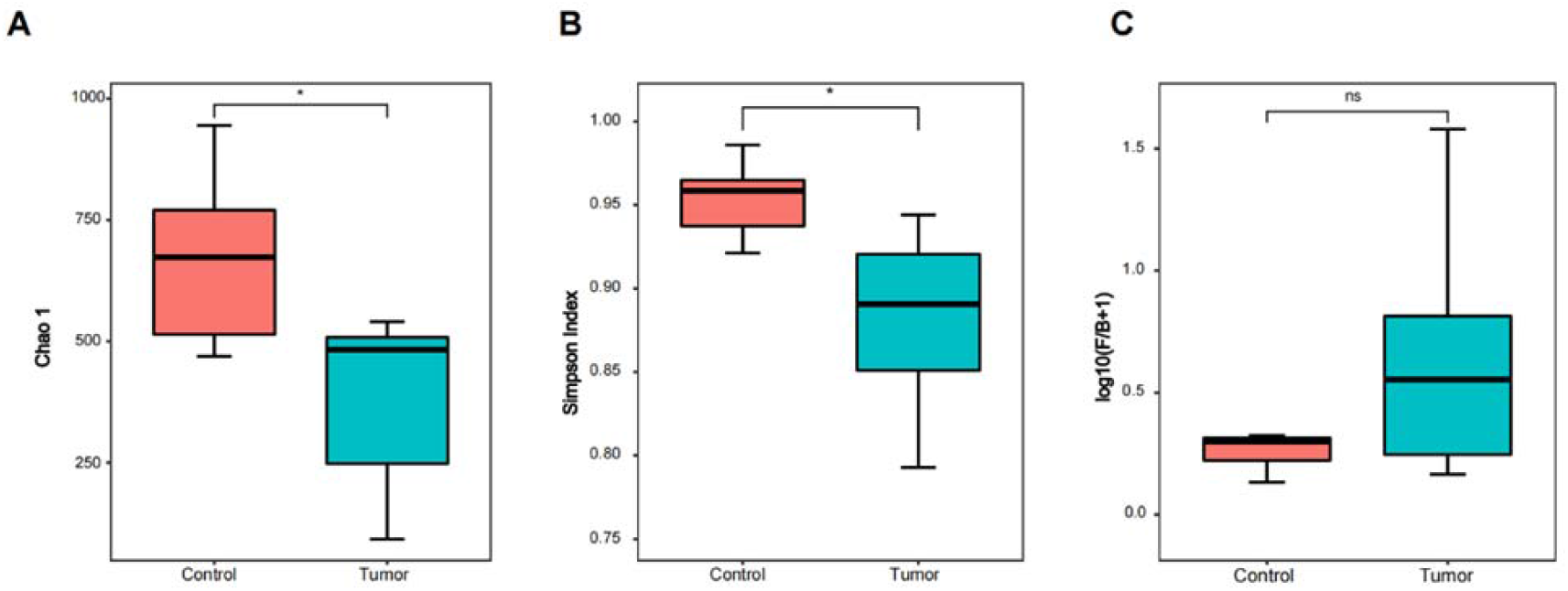
Relative bacterial richness analysis between the control and tumor groups. (A)(B) Microbial richness based on the Chao1 index and the Simpson index. *p<0.05. (C) The *Firmicutes*/*Bacteroidetes (F/B)* ratio showed no significant variation between the control and glioma model groups.

**Figure S3.**
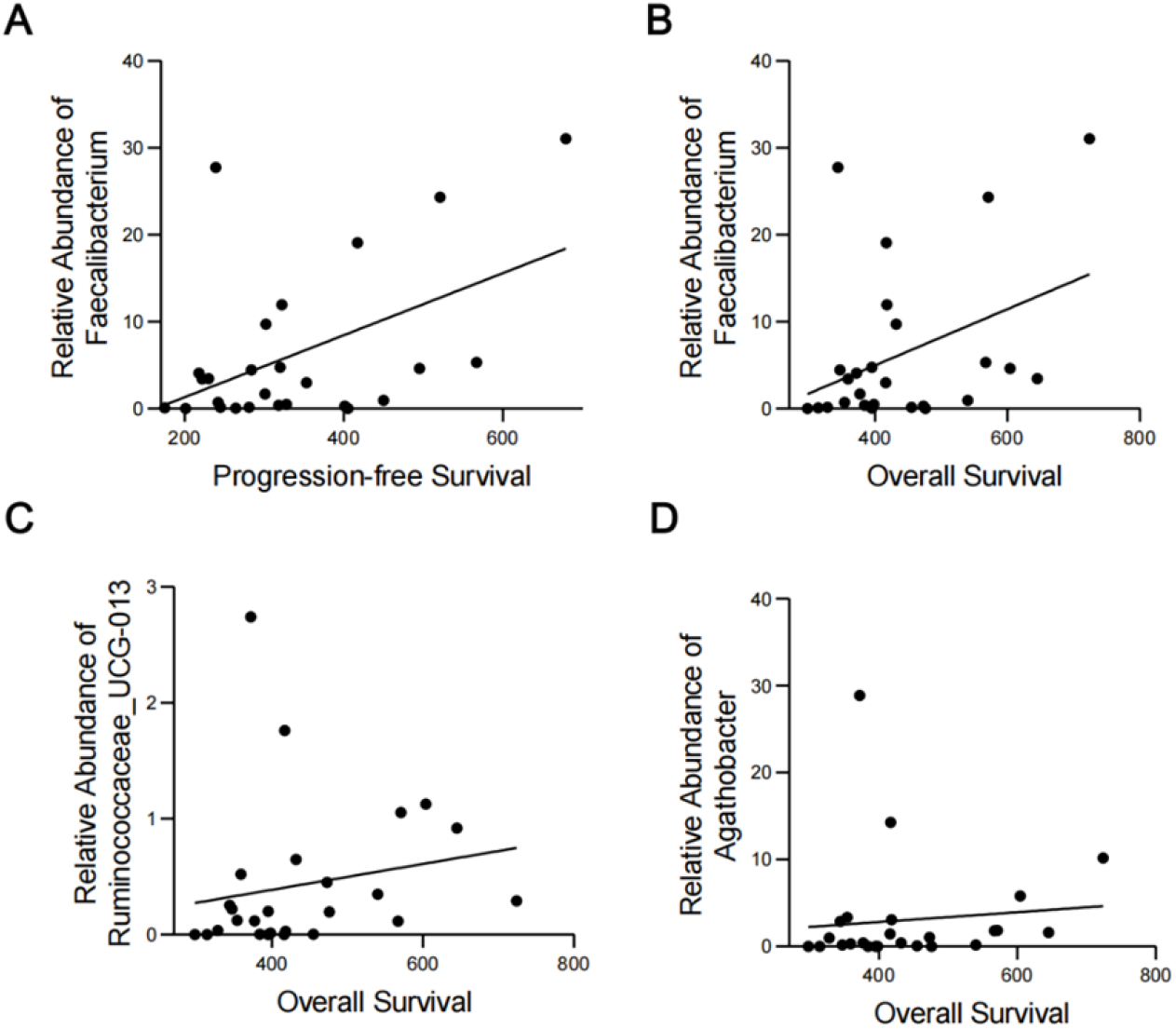
Pearman’s correlation analysis between fecal bacterial genera and PFS or OS. The correlation between Faecalibacterium and PFS (A) or OS (B). (C) The correlation between *Ruminococcaceae UCG_013* and OS. (D) The correlation between Agathobacter and OS.

